# High frequency electrical stimulation induces a long-lasting enhancement of event-related potentials but does not change the perception elicited by intra-epidermal electrical stimuli delivered to the area of secondary mechanical hyperalgesia

**DOI:** 10.1101/282335

**Authors:** José Biurrun Manresa, Ole Kæseler Andersen, André Mouraux, Emanuel N. van den Broeke

**Affiliations:** Institute of Neuroscience, Université catholique de Louvain, B-1200, Brussels, Belgium; Center for Neuroplasticity and Pain (CNAP), SMI^®^, Department of Health Science and Technology, Aalborg University, Aalborg, Denmark; Instituto de Investigación y Desarrollo en Bioingeniería y Bioinformática (IBB), CONICET-UNER, Entre Ríos, Argentina

## Abstract

High frequency electrical stimulation (HFS) of the skin induces increased pinprick sensitivity in the surrounding unconditioned skin (secondary hyperalgesia). Moreover, it has been shown that brief high intensity CO2 laser stimuli, activating both Aδ- and C-fiber nociceptors, are perceived as more intense when delivered in the area of secondary hyperalgesia. To investigate the contribution of A-fiber nociceptors to secondary hyperalgesia the present study assessed if the perception and brain responses elicited by low-intensity intra-epidermal electrical stimulation (IES), a method preferentially activating Aδ-fiber nociceptors, are increased in the area of secondary hyperalgesia. HFS was delivered to one of the two forearms of seventeen healthy volunteers. Mechanical pinprick stimulation and IES were delivered at both arms before HFS (T0), 20 minutes after HFS (T1) and 45 minutes after HFS (T2). In all participants, HFS induced an increase in pinprick perception at the HFS-treated arm, adjacent to the site of HFS. This increase was significant at both T1 and T2. HFS did not affect the percept elicited by IES, but did enhance the magnitude of the N2 wave of IES-evoked brain potentials, both at T1 and at T2. HFS induced a long-lasting enhancement of the N2 wave elicited by IES in the area of secondary hyperalgesia, indicating that HFS enhances the responsiveness of the central nervous system to nociceptive inputs conveyed by AMH-II nociceptors. However, we found no evidence that HFS affects the perception elicited by IES, which may suggest that AMH-II nociceptors do not contribute to HFS-induced secondary hyperalgesia.

## 1. INTRODUCTION

Cutaneous injury leads to increased pain sensitivity in the area of injury (primary hyperalgesia) as well as the surrounding uninjured skin (secondary hyperalgesia; Hardy et al., 1950). Secondary hyperalgesia can also be induced experimentally by activating nociceptors in an intense fashion, for example by applying high-frequency electrical stimulation (HFS) onto the skin. Indeed, HFS induces a pronounced increase in mechanical pinprick sensitivity, extending well beyond the skin area onto which HFS is applied, and lasting several hours (Klein et al., 2004; Pfau et al., 2011; Henrich et al., 2015; Van den Broeke et al., 2016b).

Whether secondary hyperalgesia evokes increased responses to thermal stimuli remains debated as studies have reported contradicting results. Indeed, whereas some studies report an increase in heat sensitivity in the area of secondary hyperalgesia induced by intradermal capsaicin injection (LaMotte et al., 1991; Serra et al., 1998; Sumikura et al., 2005) other studies do not (Simone et al., 1991; Ali et al., 1996; Geber et al., 2007).

When HFS is used to induce secondary hyperalgesia, Van den Broeke and Mouraux (2014a) have shown that brief CO2 laser stimuli heating the skin above the threshold of heat-sensitive Aδ- and C-fiber nociceptors are perceived as more intense when delivered in the area of secondary hyperalgesia (van den Broeke and Mouraux, 2014a). Brief heat stimuli can be expected to preferentially activate quickly-adapting thermonociceptors: A-fiber mechano-heat-nociceptors type II (AMH-II; Treede et al., 1998) and quickly-adapting C-fiber mechano-heat nociceptors (CMH; Meyer and Campbell, 1981; Wooten et al., 2014), and the increased heat perception following HFS could be explained by an enhancement of the responses elicited by activation of these afferents.

It has been shown that intra-epidermal electrical stimulation (IES) using a needle electrode applied against the skin can elicit responses that are exclusively related to the activation of Aδ-fiber nociceptors, provided that low stimulation intensities are used to avoid co-activation of low-threshold mechanoreceptors located more deeply in the skin. Indeed, Mouraux et al. (2010) showed that the perception and event-related brain potentials (ERPs) elicited by IES delivered at twice the absolute detection threshold are abolished in skin pre-treated with topical capsaicin to induce a reversible denervation of free nerve endings expressing the TRPV-1 receptor in the epidermal layer of the skin, while the perception and ERPs elicited by conventional transcutaneous electrical stimulation are preserved. Second, they showed that IES does not elicit neither perception nor measurable brain responses when the stimuli are delivered during an A-fiber nerve conduction block, suggesting that C-fibers, which are unaffected by the block, do not significantly contribute to the responses elicited by IES. Based on these results, it has been suggested that the responses elicited by low-intensity IES are mainly related to the activation of capsaicin-sensitive Aδ-fiber nociceptors, which includes the AMH type II (Mouraux et al. 2010; Liang et al., 2016). The aim of the present study was to test whether the responses elicited by IES are increased in the area of secondary hyperalgesia induced by HFS, and, to examine whether this enhancement correlates with secondary mechanical hyperalgesia.

## 2. MATERIALS AND METHODS

### 2.1 Participants

Seventeen healthy volunteers took part in the experiment (10 men and 7 women; aged 26.8 ± 3.3 years [mean ± sd]). The experiment was conducted in accordance with the declaration of Helsinki and approval for the experiment was obtained from the local Ethical Committee of Aalborg Kommune (VN 2015-0038). All participants signed an informed consent form and received financial compensation for their participation.

### 2.2 Experimental procedure

The design of the experiment is summarized in Figure 1. During the experiment, participants were comfortably seated in an adjustable chair with their arms in supine position resting on a table in front of them. Mechanical pinprick stimuli and IES were administered to both arms before (T0), 20 minutes after (T1) and 45 minutes after (T2) transcutaneous HFS of the volar site of one of the forearms.

**Figure 1.**
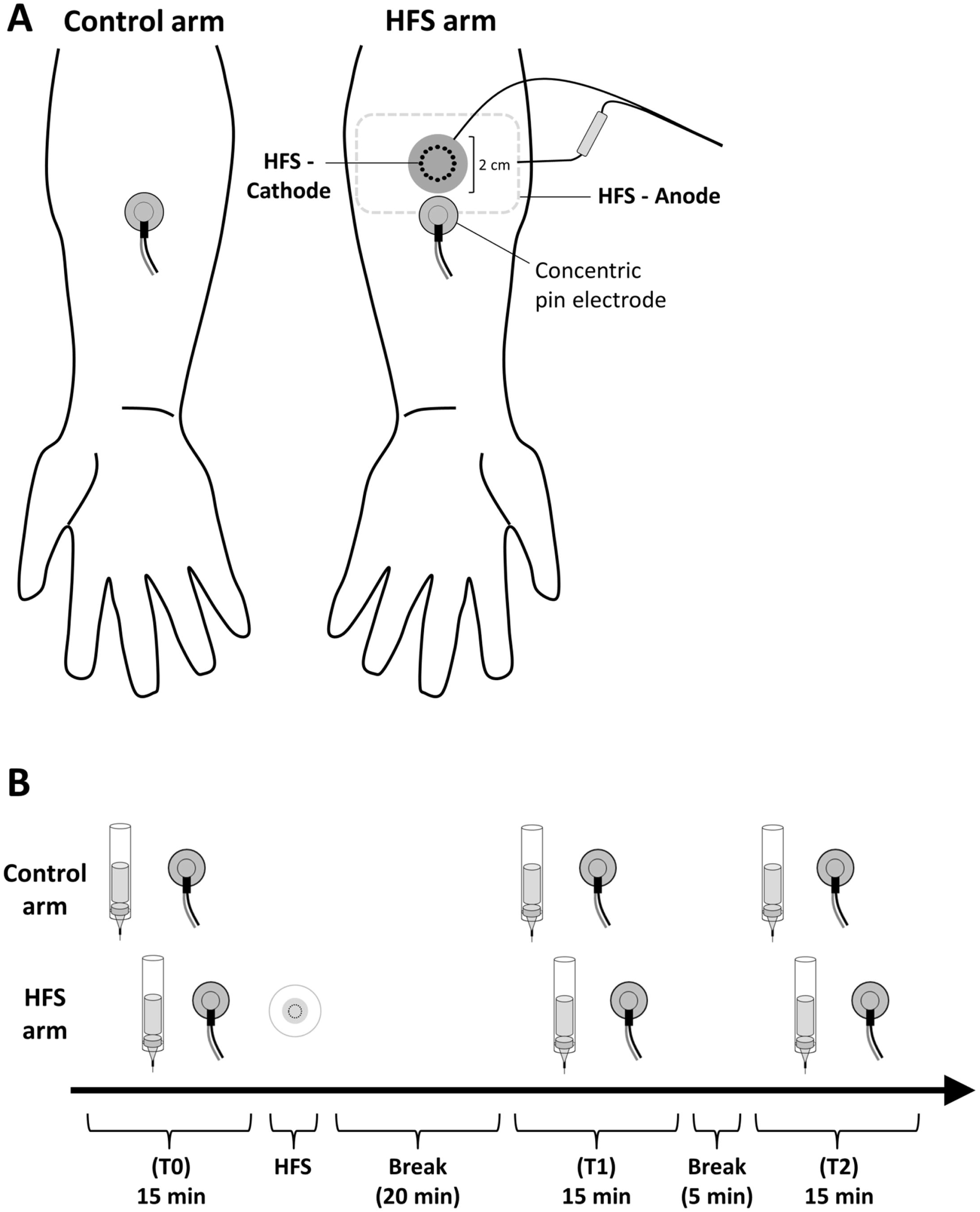
**A.** Experimental set up. High frequency electrical stimulation of the skin (HFS) was applied on either the left or right volar forearm. Intra-epidermal electrical stimuli (IES) were delivered to the left and right volar forearm using a concentric pin electrode, placed adjacent to the HFS electrode. Mechanical pinprick stimuli were applied to the skin surrounding the IES electrode. **B.** Time-line of the experiment. The effects of HFS on the perception to mechanical pinprick stimuli and the perception and brain responses elicited by IES were assessed at 3 different time points: before HFS (T0), 20 min after HFS (T1), and 45 min after HFS (T2).

### 2.3 High frequency electrical conditioning stimulation (HFS)

HFS was applied to the volar site of one of the two forearms, as described in Van den Broeke et al. (2016a; 2017). To avoid any confounding effect of handedness, the arm onto which HFS was applied (dominant vs. non-dominant) was counterbalanced across participants. Handedness was assessed using the Flinders Handedness Survey (Nicholls et al., 2013). The HFS protocol consisted of five trains of electrical pulses delivered at 100 Hz (pulse width: 2 ms). Each train lasted 1 s, and the time interval between the onsets of each train was 10 s. The electrical pulses were triggered by custom-made software (“Mr. Kick”, Aalborg University, Aalborg, Denmark), generated by a computer-controlled constant-current electrical stimulator (Noxitest IES 230, Aalborg, Denmark), and delivered to the skin using a custom-made electrode built at the Centre for Sensory-Motor Interaction (SMI^®^, Aalborg University, Denmark). The cathode consisted of 16 blunt stainless-steel pins with a diameter of 0.2 mm protruding 1 mm from the base. The 16 pins are placed in a circle with a diameter of 10 mm. The anode consisted of a flexible electrode pad (50 mm × 90 mm, type Synapse, Ambu A/S, Denmark) that was attached to the dorsum of the arm. The intensity of HFS was individually adjusted to 20 times the detection threshold to a single electrical pulse delivered using a DS5 constant-current stimulator (Digitimer, UK). The thresholds were determined using an automatic staircase procedure: stimuli were initially delivered at an intensity of 100 μA and increasing in 50 μA steps until the stimulus was perceived by the subject, signalled by pushing a button as fast as possible (within a time window of 1.6 s). Afterwards, the intensity was decreased in steps of 10 μA until the stimulus was no longer detected. Then, the intensity was increased in steps of 10 μA until the stimulus was detected again. Three ascending and three descending staircases were applied, and the detection threshold was defined as the average intensity of the last two peaks and troughs. The inter-stimulus interval was random between 5-8 s.

### 2.4 Mechanical pinprick stimulation

To assess heterotopic changes in pinprick sensitivity, a calibrated pinprick stimulator exerting a normal force of 128 mN (“The Pin Prick”, MRC Systems, Heidelberg, Germany) was applied perpendicular to the skin. A total of three pinprick stimuli were applied adjacent to the concentric pin electrode on the HFS-conditioned arm and on the corresponding location of the contralateral control arm. To avoid sensitization of the stimulated skin by the mechanical stimuli, the target of each pinprick stimulus was slightly displaced after each stimulus. Participants were asked to report the mean perceived intensity elicited by the three pinprick stimuli, on a numerical rating scale (NRS) ranging from 0 (no perception) to 100 (maximal pain), with 50 representing the transition from non-painful to painful domains of sensation.

### 2.5 Intra-epidermal electrical stimulation (IES)

IES consisted of two succeeding constant-current square-wave pulses (“double pulse”), with each a duration of 0.5 ms, separated by a 10 ms inter-pulse interval (Mouraux et al., 2010). The stimuli were generated by a constant-current stimulator (DS5, Digitimer, UK) and delivered using a stainless steel concentric bipolar needle electrode developed by Inui et al. (Inui et al., 2002). The electrode consists of a needle cathode (length: 0.1 mm, Ø: 0.2 mm) and a surrounding cylindrical anode (Ø: 1.4 mm). By gently pressing the electrode onto the skin, the needle electrode was inserted into the epidermis. The electrode was then fixated to the skin with self-adhesive tape. The intensity of IES was individually adjusted to twice the detection threshold to a double pulse. The thresholds were established using the same staircase procedure that was used to assess the detection threshold for HFS. At each time point (T0, T1 and T2) and each arm (HFS and control arm) a block of thirty double pulse stimuli were delivered using a random inter-stimulus interval ranging from 8 to 10 s. During IES, subjects were instructed to push a button, held in the hand opposite to the arm being stimulated, as fast as possible when they perceived a stimulus (detection of reaction time). To assess the changes in the intensity of the percept elicited by IES, participants were asked to rate directly after each block of thirty stimuli the intensity on the same NRS that was used to rate the mechanical pinprick stimuli.

### 2.6 EEG recording

The electroencephalogram (EEG) was recorded using nine active electrodes mounted in an elastic electrode-cap (g.SCARABEO, g.tec, Medical Engineering GMBH, Austria) and placed according to the international 10-20 system (electrode locations: FP1, AFz, Fz, Cz, C1, C2, C3, C4 and Pz). Participants were instructed to keep their gaze fixed on a black cross displayed at approximately 2.0 m distance (∼30° below eye level) and to sit as still as possible. The EEG signals were amplified and digitized using a sampling rate of 1200 Hz and a left earlobe (A1) reference (g.Hlamp, g.tec, Medical Engineering GMBH, Austria). The ground electrode was placed at position AFz. Electrode impedances were kept below 20 kΩ as assessed by the g.tec EEG system.

### 2.7 Data analysis

#### EEG pre-processing

The EEG signals were analysed offline using Letswave 6.0 (www.nocions.org/letswave). After applying a 0.3 - 30 Hz band pass zero-phase Butterworth filter (4th order) to the continuous EEG recordings, the signals were segmented into epochs extending from −500 to +1500 ms relative to stimulus onset. Epochs contaminated by eye movements or eye blinks were corrected using the Gratton and Coles method (Gratton et al., 1983). After applying a baseline correction (reference interval: −500 to 0 ms), epochs with amplitude values exceeding ± 100 μV were rejected as these were likely to be contaminated by artefacts. Finally, separate average waveforms were computed for each participant, time point (T0, T1 and T2) and stimulation site (HFS and control). To determine the latency window of the ERPs, the grand average global field power (GFP) including all conditions and all participants was calculated (Van den Broeke and Mouraux, 2014a). Two distinct peaks (N2 and P2) were identified in the grand average global field power (Fig. 6A). The N2 was defined as the most negative peak within the time interval extending from 130 to 230 ms, and the P2 was defined as the most positive peak within the time interval extending from 280 to 500 ms. Peak amplitudes were expressed relative to baseline. Peak latencies were expressed relative to stimulus onset.

**Figure 2.**
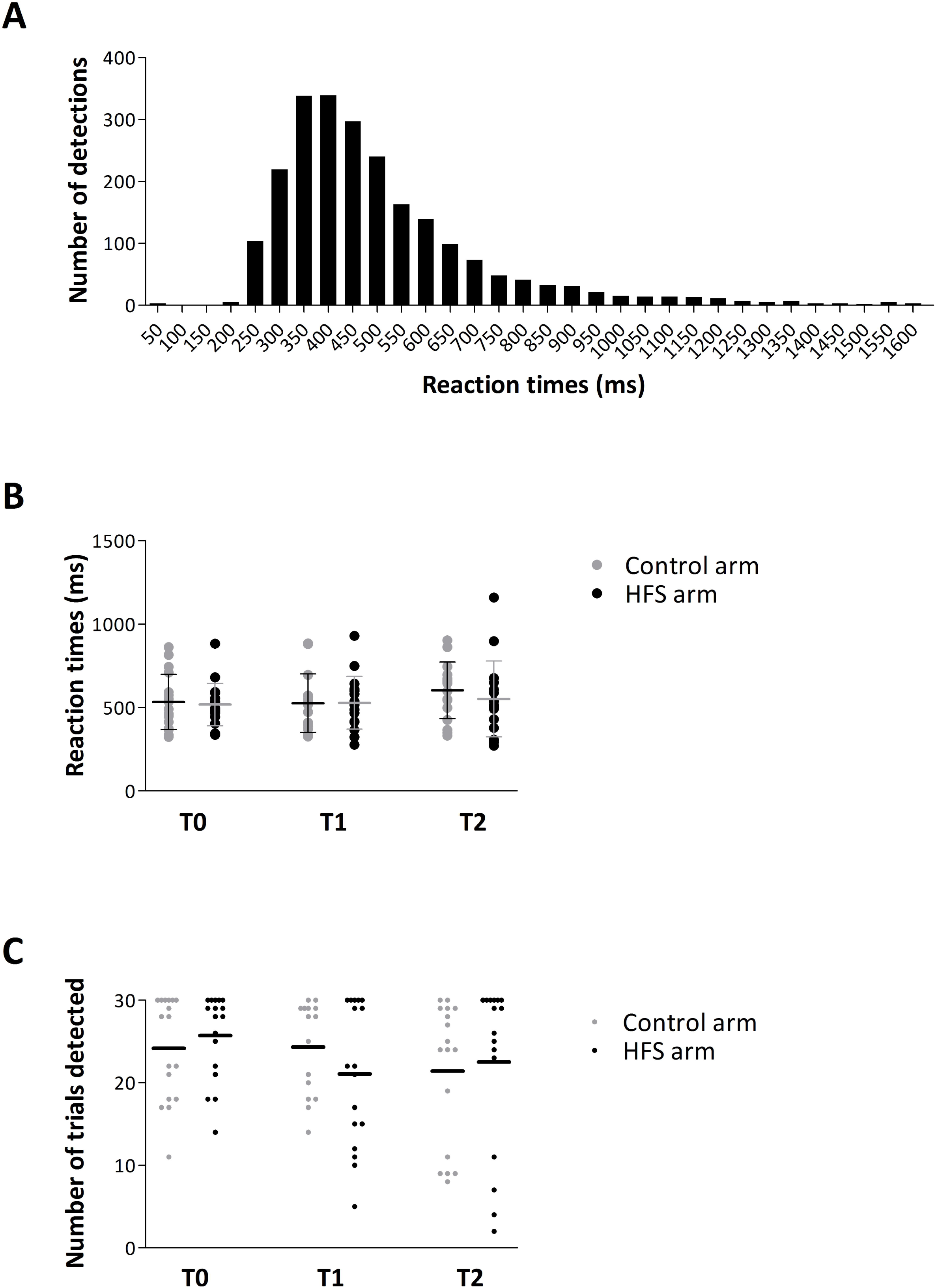
**A.** Distribution of reaction times to intra-epidermal electrical stimuli delivered at all time-points (T0, T1 and T2) and arms (control and HFS). Note that the distribution peaks at latencies compatible with the conduction velocity of Aδ fibers. **B.** Individual and group-level average (±SD) reaction times for each time-point and arm. **C.** Number of detected trials for each participant at each time-point and arm. Horizontal bar shows the group-level average number of detected trials.

**Figure 3.**
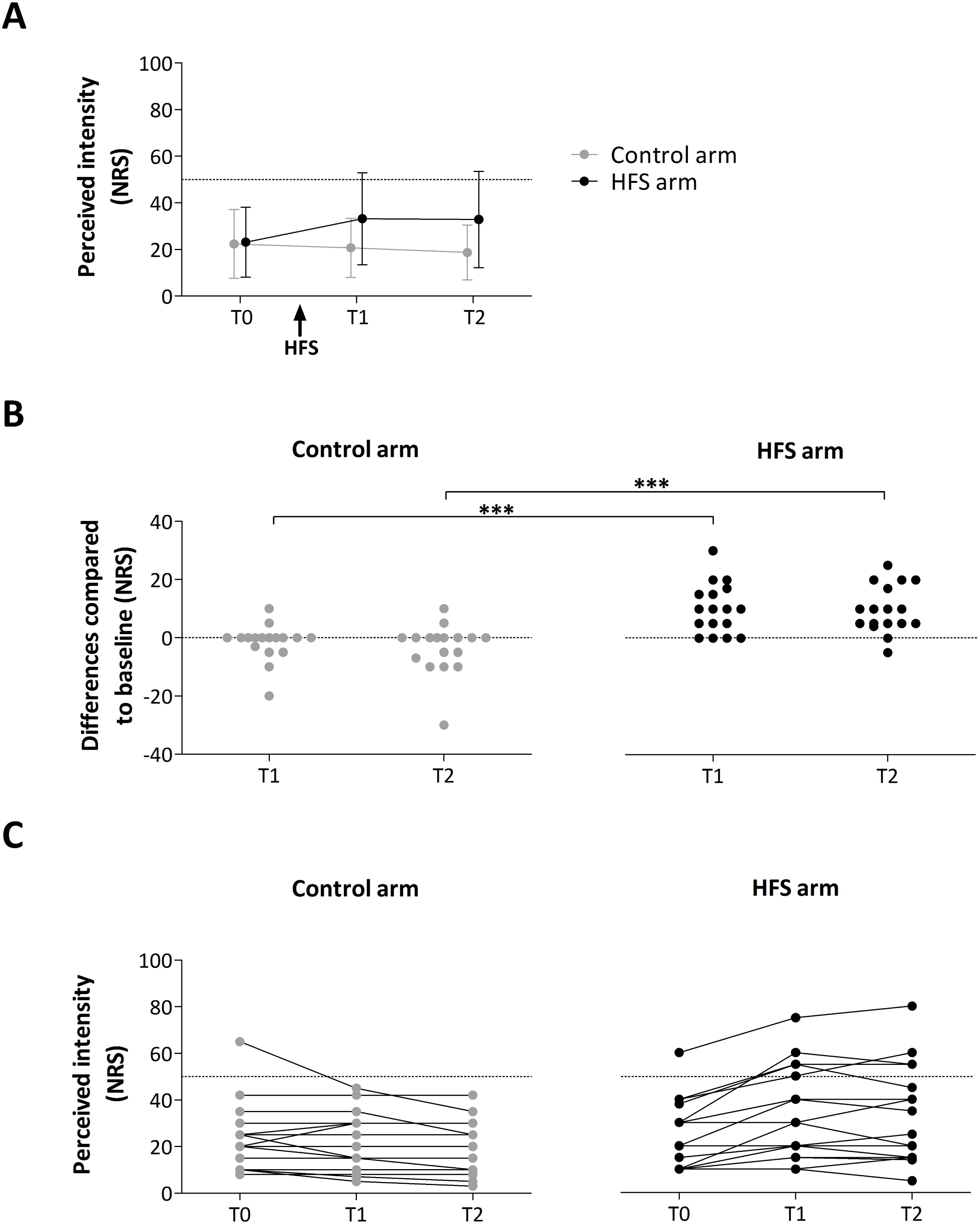
**A.** Group-level average (± SD) intensity of perception elicited by mechanical pinprick stimuli. **B.** Individual difference ratings (post-pre) at time point T1 and T2 for control and HFS arm. Asterisks show the significant enhancement of pinprick perception after HFS at the HFS treated arm. *** = *p*<.001. **C.** Individual ratings for both arms (control and HFS) at all time points (T0, T1 and T2).

**Figure 4.**
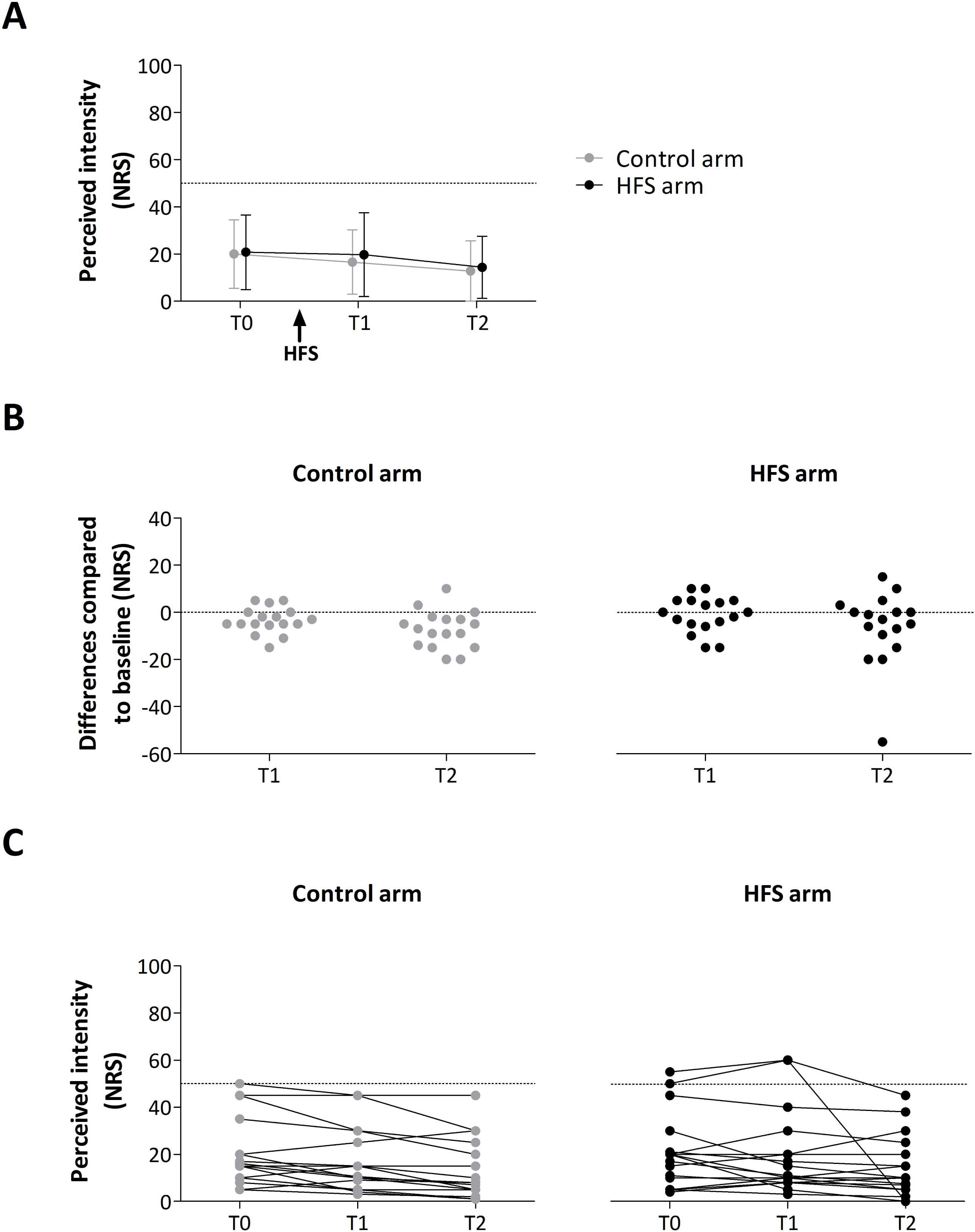
**A.** Group-level average (± SD) intensity of perception elicited by intra-epidermal electrical stimuli. **B.** Individual difference ratings (post-pre) at time point T1 and T2 for control and HFS arm. Note that the distribution of the data points at the HFS arm is not bimodal which argues against the existence of two different subgroups (“responders” vs “non-responders”). **C.** Individual ratings for both arms (control and HFS) at all time points (T0, T1 and T2).

**Figure 5.**
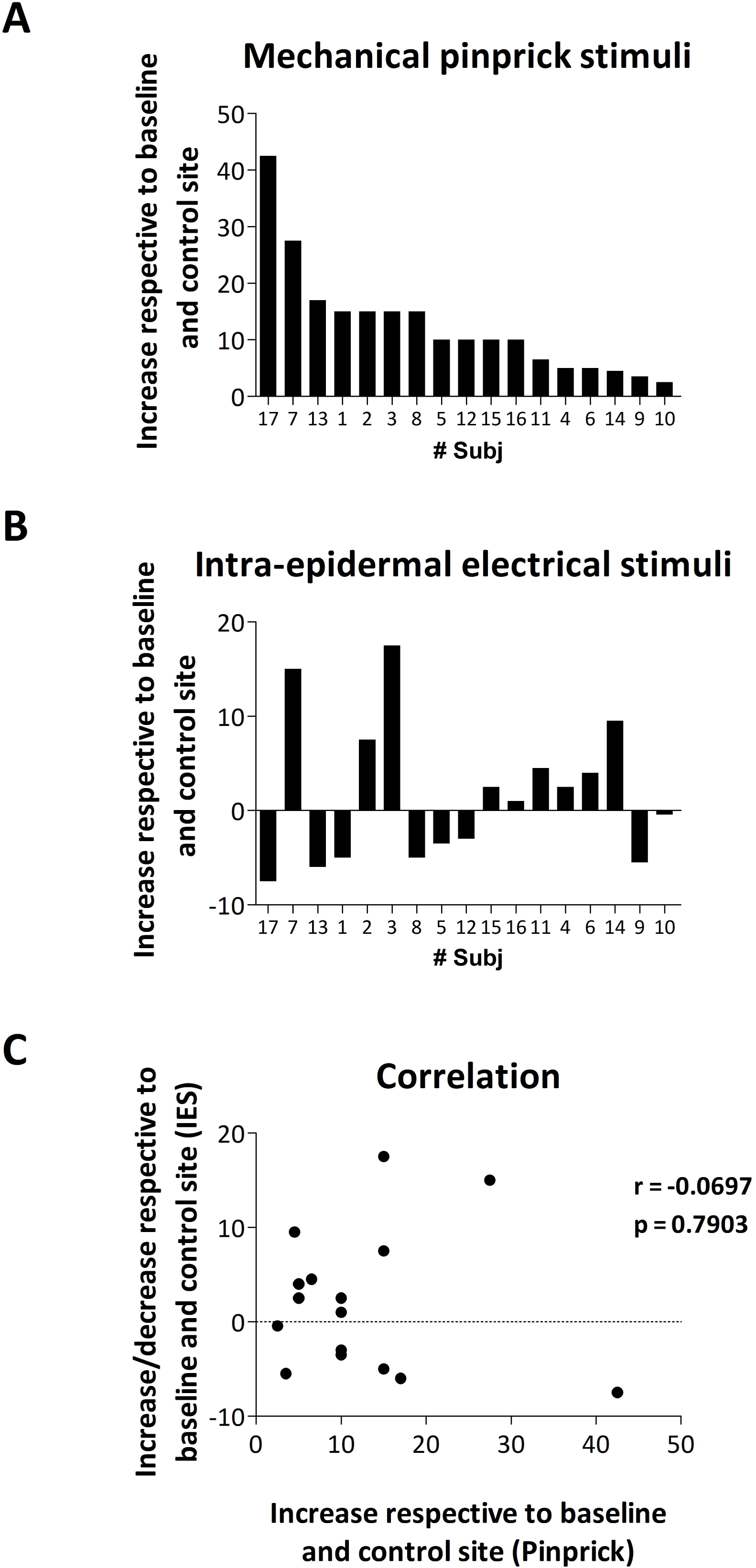
**A.** Increase in pinprick perception after HFS (respective to baseline and control site) for each participant and ordered according to size, from large to small. **B.** Increase or decrease in the perception elicited by intra-epidermal electrical stimulation after HFS (respective to baseline and control site) for every participant and according to the order in A. **C.** Lack of correlation between the increase in pinprick perception and the increase/decrease in the perception to intra-epidermal electrical stimulation.

**Figure 6.**
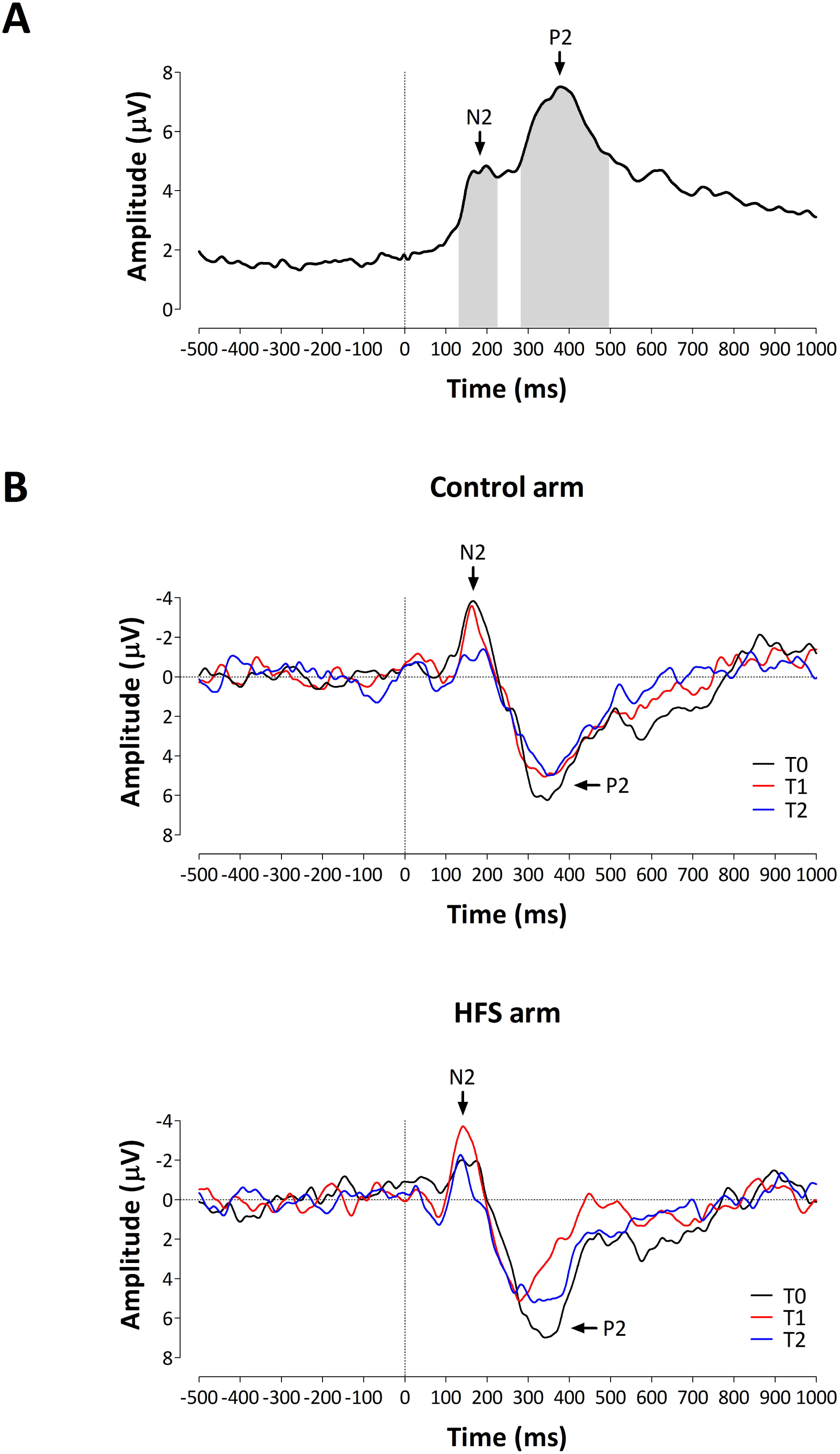
**A.** Grand-average global field power of the event-related potentials elicited by intra-epidermal electrical stimulation. The global field power is calculated across all electrodes, conditions and participants. Two ERP peaks can be identified. First, a negative peak appearing between 130 and 230 ms, labelled N2. Second, a positive peak appearing between 280 and 500 ms, labelled P2. The grey squares represent the time window that was used for identifying the N2 and P2 peaks in individual waveforms. **B.** Group-level average event-related potentials (Cz) elicited by intra-epidermal electrical stimulation before and after applying HFS from both arms.

### 2.8 Statistical analysis

A two-way repeated measures analysis of variance (ANOVA) was performed to assess differences in intensity of perception elicited by the mechanical pinprick stimuli and IES, reaction times, detection rates, and N2 and P2 amplitudes and latencies. For reaction times and detection rates, the average values across 30 trials were used as dependent variable. *Time* (T0, T1 and T2) and *arm* (control vs. HFS arm) were used as within-subject factors. The assumption of sphericity was tested using Mauchly’s test. In case the data violated the assumption of sphericity, F-values were corrected using the Greenhouse-Geisser procedure (denoted F_G-G_). The effects of HFS were further assessed using planned contrasts (“simple” method). The level of significance was set at *p* < 0.05 (two-sided). To investigate the relationship between the changes in perceived pinprick intensity and IES after HFS, the Pearson’s correlation coefficient was calculated between the variation in perceived pinprick intensity and the perceived intensity elicited by IES. Values are stated as mean ± standard deviation unless stated otherwise. The statistical analyses were conducted using SPSS 18 (SPSS Inc., Chicago, IL, USA).

## 3. RESULTS

### 3.1 Detection thresholds

The detection threshold to a single pulse of HFS was 170 ±50 µA (Min: 75 μA, Max: 260 μA). The detection thresholds to a double pulse of IES were 69 ±25 μA (Min: 37.5 μA, Max: 113 μA) for the control arm and 83 ±35 μA (Min: 40 μA, Max: 155 μA) for the HFS arm. This difference was not significant (t (16) = 2.058, *p* = 0.0562).

### 3.2 Reaction times and detection rates

Average reaction times to IES before and after applying HFS (T0, T1 and T2) at both arms are depicted in Figure 2. For two subjects, the reaction times at T1 for the control side were not saved due to technical problems, whereas one subject did not detect any of the IES stimuli delivered at the HFS arm at T2. The ANOVA performed on the reaction times of the remaining 14 participants did not reveal a significant main effect of time (F-(2,26) = 1.813, *p* = 0.183, partial η^2^ = 0.122), main effect of arm (F-(1,13) = 0.022, *p* = 0.885, partial η^2^ = 0.002) or time x arm interaction (F_G-G_-(1.268,16.485) = 0.512, *p* = 0.527, partial η^2^ = 0.038). Figure 2C shows the number of IES that were detected at each time-point for each participant and arm. The ANOVA performed on the detection rates of the 15 participants did not reveal a significant main effect of time (F_G-G_-(1.376,19.266) = 3.336, *p* = 0.072, partial η^2^ = 0.192), or arm (F-(1,14) = 0.943, *p* = 0.348, partial η^2^ = 0.063) or time x arm interaction (F-(2,28) = 1.258, *p* = 0.300, partial η^2^ = 0.082).

### 3.3 Intensity of perception

#### 3.3.1 Mechanical pinprick stimulation

HFS induced a clear increase in mechanical pinprick sensitivity (Figure 3). This was confirmed by the two-way repeated measures ANOVA that showed a significant main effect of time (F-(2,32) = 5.719, *p* = 0.008, partial η^2^ = 0.263), a significant main effect of arm (F-(1,16) = 19.367, *p* < 0.001, partial η^2^ = 0.548) and a significant time x arm interaction (F_G-G_-(1.481,23.698) = 20.744, *p* < 0.001, partial η^2^ = 0.565). The univariate within-subject contrasts revealed that the mechanical pinprick ratings were significantly enhanced at the conditioned arm after HFS at T1 (F-(1,16) = 26.986, *p* < 0.001, partial η^2^ = 0.628) and T2 (F-(1,16) = 23.012, *p* < 0.001, partial η^2^ = 0.590).

#### 3.3.2 Intra-epidermal electrical stimulation

The intensity of perception elicited by IES delivered to the control and HFS-conditioned arm before and after HFS (T0, T1 and T2) is shown in Figure 4. The ANOVA revealed a significant main effect of time (F_G-G_-(1.334, 21.352) = 5.652, *p* < 0.019, partial η^2^ = 0.590). The univariate within-subject contrasts revealed that the IES ratings at T2 were significantly lower compared to T0 (F-(1, 16) = 6.749, *p* = 0.019, partial η^2^ = .297). No significant main effect of arm (F-(1, 16) = 2.960, *p* = 0.105, partial η^2^ = .156) or time × arm interaction (F_G-G_-(1.459, 23.345) = 0.372, *p* = 0.627, partial η^2^ = .023) was observed.

#### 3.3.3 Correlation between mechanical pinprick and IES ratings

Figure 5 shows the variation in ratings after HFS respective to baseline and control site for mechanical pinprick stimuli and IES. T1 and T2 post-measurements were merged. Figure 5C shows that there was no linear correlation between the two modalities regarding the increase in ratings after HFS.

### 3.4 Event-related brain potentials elicited by IES

The group-level average event-related potentials elicited by IES before (T0) and after applying HFS (T1 and T2), for each arm (control and HFS) are shown in Figure 6B.

#### 3.4.1 N2 magnitude

The magnitude of the vertex N2 wave elicited by IES delivered to the control and HFS-conditioned arm before (T0) and after (T1, T2) conditioning is shown in Figure 7A and Table 1. The ANOVA performed on the magnitude of the N2 wave showed no significant main effect of time (F-(2,32) = 2.296, *p* = 0.117, partial η^2^ = 0.125), no significant main effect of arm (F-(1,16) = 0.180, *p* = 0.677, partial η^2^ = 0.011), but a significant time × arm interaction (F-(2,32) = 5.501, *p* = 0.009, partial η^2^ = .256). The univariate within-subject contrasts revealed that the magnitude of the N2 wave was significantly enhanced at the conditioned arm after HFS at T2 (F-(1,16) = 14.323, *p* = 0.002, partial η^2^ = .472) and almost at T1 (F-(1,16) = 4.045, *p* = 0.061, partial η^2^ = .202; Figure 7C).

**Table 1.** Group-level mean (± SD) amplitude and latency of the ERP N2 and P2 elicited by intra-epidermal electrical stimuli before and after HFS at both arms.

|  |  | N2 wave |  |  |  |  |  | P2 wave |  |  |  |  |  |
| --- | --- | --- | --- | --- | --- | --- | --- | --- | --- | --- | --- | --- | --- |
| | | Amplitude (μV) | | | Latency (ms) | | | Amplitude (μV) | | | Latency (ms) | | |
|  |  | *T0* | *T1* | *T2* | *T0* | *T1* | *T2* | *T0* | *T1* | *T2* | *T0* | *T1* | *T2* |
| **Control** | Mean | -6.2 | -5.5 | -3.4 | 180 | 186 | 181 | 8.4 | 8.1 | 6.6 | 414 | 394 | 397 |
|  | SD | 5.0 | 5.5 | 3.4 | 38 | 25 | 29 | 4.8 | 5.3 | 5.8 | 62 | 61 | 67 |
| **HFS** | Mean | -4.8 | -6.6 | -5.3 | 181 | 185 | 178 | 9.1 | 7.9 | 8.0 | 376 | 355 | 361 |
|  | SD | 4.8 | 7.1 | 5.3 | 37 | 33 | 37 | 6.6 | 6.8 | 7.1 | 41 | 66 | 56 |

**Figure 7.**
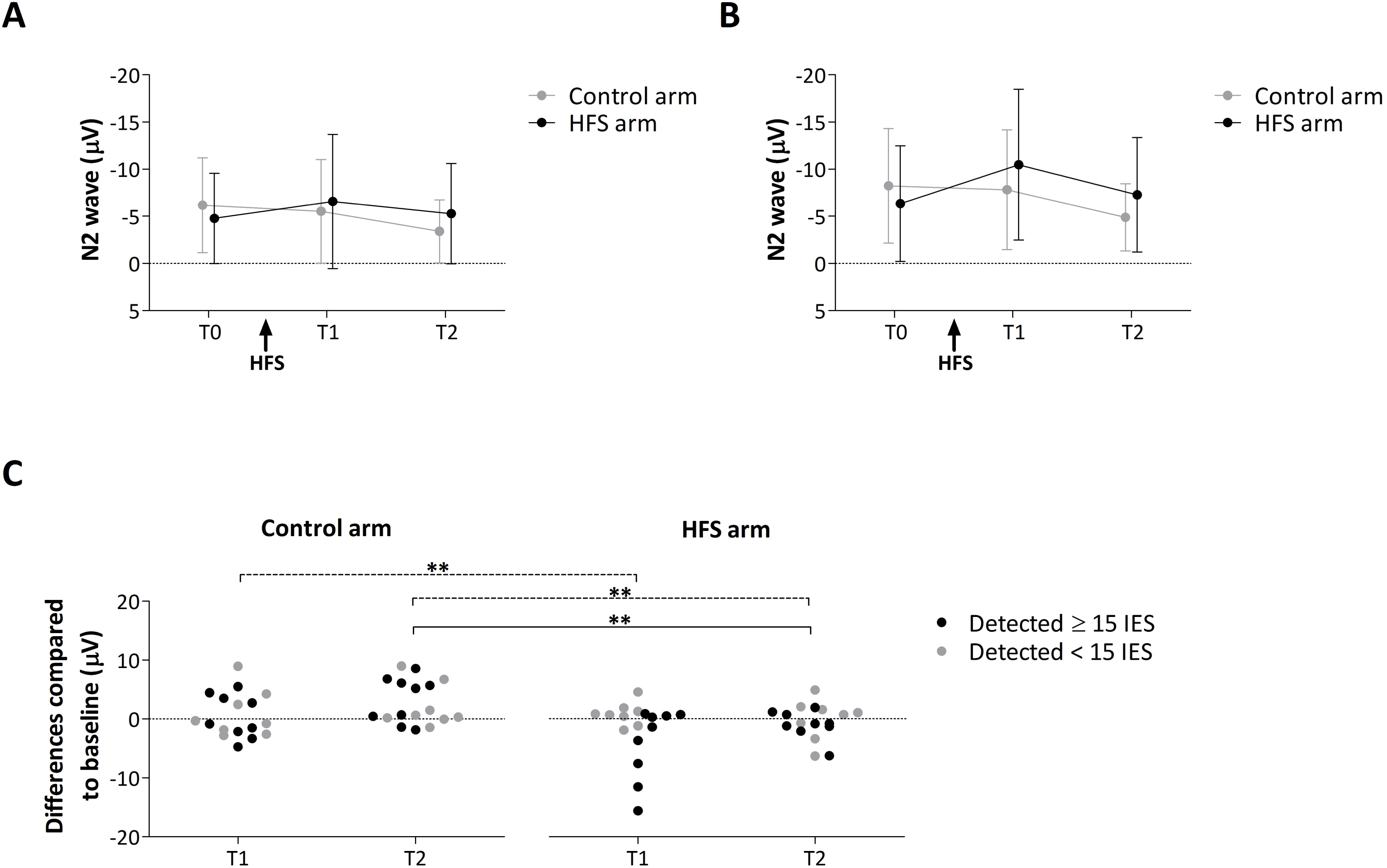
**A.** Group-level average (± SD) magnitude of the N2 wave elicited by intra-epidermal electrical stimuli before and after HFS at the two arms. **B.** Group-level average (± SD) magnitude of the N2 wave for the 9 participants that detected the stimulus in more than half of the trials for each condition. **C.** Individual difference ratings (post-pre) at time point T1 and T2 for control and HFS arm. Black dots represent the participants that detected at least half of the stimuli in every condition, whereas the grey dots represent the participants that detected less than half of the stimuli. A negative or positive value indicates an enhancement or decrease of the N2 wave, respectively. Asterisks show the significant enhancement of the N2 wave after HFS at the HFS treated arm. ** = *p*<.01. Solid line refers to the main analysis (all participants) whereas the dashed lines refer to the additional analysis (participants that detected at least half of the stimuli).

#### 3.4.2 N2 latency

The two-way repeated measures ANOVA performed on the N2 latency did not reveal a significant main effect of time (F_G-G_-(1.376,22.011) = 0.371, *p* = 0.616, partial η^2^ = 0.023), main effect of arm (F-(1,16) = 0.024, *p* = 0.879, partial η^2^ = 0.001) or time x arm interaction (F-(2,32) = 0.044, *p* = 0.957, partial η^2^ = 0.003).

#### 3.4.3 P2 magnitude

The two-way repeated measures ANOVA performed on the P2 magnitude did not reveal a significant main effect of time (F-(2,32) = 2.227, *p* = 0.124, partial η^2^ = 0.122), main effect of arm (F-(1,16) = 0.357, *p* = 0.558, partial η^2^ = 0.022) or time x arm interaction (F-(2,32) = 0.348, *p* = 0.709, partial η^2^ = 0.021).

#### 3.4.4 P2 latency

The two-way repeated measures ANOVA performed on the P2 latency did not reveal a significant main effect of time (F-(2,32) = 1.838, *p* = 0.175, partial η^2^ = 0.103). However, the ANOVA did reveal a significant main effect of arm (F-(1,16) = 10.235, *p* = 0.006, partial η^2^ = 0.390), in which P2 latencies for the control arm were, on average, longer than for the HFS arm. No significant time x arm interaction (F-(2,32) = 0.022, *p* = 0.978, partial η^2^ = 0.001) was observed.

#### 3.4.5 Correlation between mechanical pinprick ratings and N2 wave elicited by IES

Figure 8 shows the lack of correlation between the increase in pinprick ratings and the increase in magnitude of the vertex N2 wave elicited by IES.

**Figure 8.**
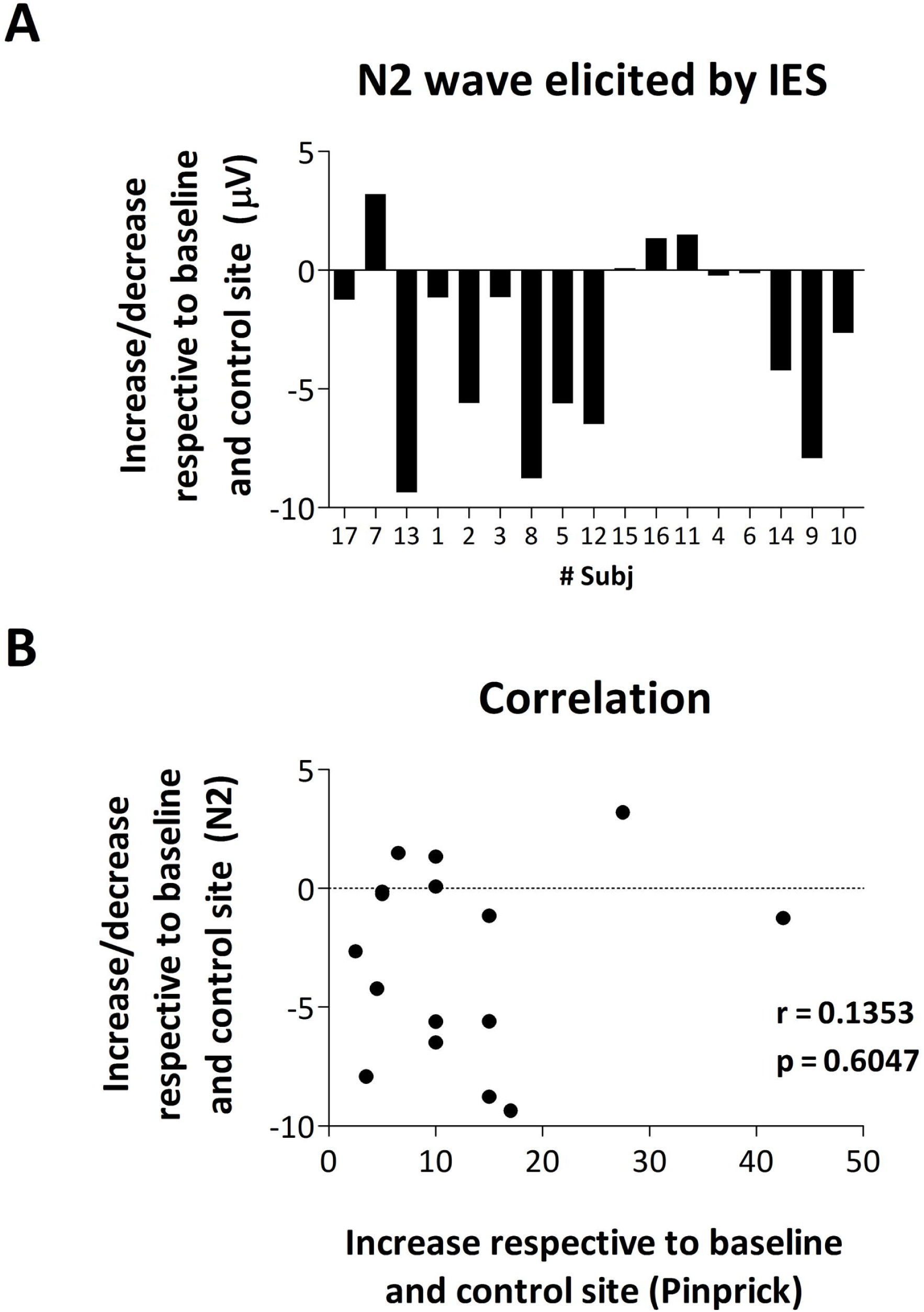
**A.** Increase or decrease in magnitude of the N2 wave of IES-evoked brain potentials after HFS (respective to baseline and control site) for every participant. **B.** Lack of correlation between the increase in pinprick perception and the change in magnitude of the N2 wave of IES-evoked brain potentials.

### 3.5 Additional analysis

Several participants failed to detect the IES in more than half of the trials in at least one of the six recording sessions (Fig. 2C: 8/17 participants). Therefore, we performed an additional analysis of the N2 wave magnitude taking into account only the participants that detected at least half of the total number of trials in every condition (N=9). In accordance with the main analysis, the ANOVA revealed a significant time x arm interaction (F-(2, 16) = 10.213, *p* = 0.001, partial η^2^ = 0.561). The univariate within-subject contrasts revealed that the magnitude of the N2 wave was significantly enhanced at the conditioned arm at both T1 (F-(1, 8) = 13.785, *p* = 0.006, partial η^2^ = 0.633) and T2 (F-(1, 8) = 13.125, *p* = 0.007, partial η^2^ = 0.621). The N2 magnitude of each time-point (T0, T1 and T2) and arm (control and HFS) are shown in Figure 7B. The group-level ERPs are shown in Figure S1 of the *Supporting information*.

## 4. DISCUSSION

The present study shows that HFS does not increase the perception elicited by IES delivered in the area of secondary mechanical hyperalgesia. However, HFS does enhance the magnitude of the vertex N2 wave elicited by IES. Such as the after-effect of HFS on pinprick sensitivity, the effect of HFS on the N2 wave is long-lasting, both being present both 20 minutes (T1) and 45 minutes (T2) after applying HFS. In contrast, no significant enhancement of the P2 wave was observed.

### 4.1 HFS does not increase the perception elicited by IES

The perception elicited by IES was not changed by HFS. This suggests that AMH-II nociceptors do not contribute to HFS-induced secondary hyperalgesia. This observation is in contrast with the results of Liang et al. (2016) examining the after-effects of intradermal capsaicin on the responses to IES. By dividing their sample into “responders” and “non-responders” based on the amount of pinprick hypersensitivity observed after capsaicin injection, they found that the percept elicited by IES was increased in “responders” (N=6), whereas it was decreased in “non-responders” (N=6). Their results suggested a relationship between the amount of pinprick hypersensitivity and the changes in the perception of IES. In the present study, although we recruited a larger number of participants (n=17), we did not observe such a relationship. Despite the fact that all our participants exhibited an increase in pinprick sensitivity after HFS, i.e. all were “responders”, there was no significant increase in the percept elicited by IES, and no correlation between the magnitude of the post-HFS increase in pinprick sensitivity and the post-HFS change in the percept elicited by IES. These findings could suggest that capsaicin-induced secondary hyperalgesia is different from HFS-induced secondary hyperalgesia. A first important difference between capsaicin injection and HFS is that capsaicin selectively activates nociceptors expressing the TRPV-1 receptor, while HFS bypasses receptor transduction and activates all primary afferents indistinctly. A second difference is that the duration of conditioning stimulation is shorter for HFS than for capsaicin injection. Indeed, while HFS activates nociceptive afferents for only five seconds, injected capsaicin can be expected to generate activity in capsaicin-sensitive afferents during several minutes.

However, it is important to be cautious when comparing directly the results of Liang et al. with the present study. First, because secondary hyperalgesia was induced at different body sites (hand dorsum vs. volar forearm). Second, because the time at which IES stimulation was delivered relative to sensitization was different. While Liang et al. delivered IES directly after the sensation induced by capsaicin injection had disappeared, we delivered IES 20 and 45 minutes after HFS.

### 4.2 HFS enhances the ERP vertex negativity elicited by IES

The magnitude of the N2 wave elicited by IES was significantly enhanced after HFS at the HFS-treated arm. Like the effect of HFS on pinprick sensitivity, the effect of HFS on the magnitude of the N2 wave of IES-evoked potentials was long-lasting. Our results show that HFS enhances the responsiveness of the central nervous system to nociceptive inputs conveyed by AMH type II. Liang et al. (2016) also observed an enhancement of the N2 wave of IES-evoked potentials in a small subgroup of subjects that developed secondary mechanical hyperalgesia after intradermal capsaicin injection.

The dissociation between the IES-evoked perception and IES-evoked N2 wave suggests that the two phenomena are not necessarily related and/or reflect different processes. Dissociations between perception and the magnitude of nociceptive ERPs have been reported in previous studies (Legrain et al., 2011). Based on these observations it has been hypothesized that these brain responses reflect a system that is involved in detecting, orienting attention towards and reacting to the occurrence of salient sensory events (Legrain et al., 2011).

A previous study found that HFS also exerts a long-lasting effect on the ERPs elicited by non-nociceptive vibrotactile stimuli selectively activating fast-adapting low-threshold mechanoreceptors of the lemniscal system (Van den Broeke and Mouraux, 2014a). In that study, the authors observed a significant enhancement of the vertex N wave elicited by short-lasting vibrations delivered to the area of secondary hyperalgesia. As in the present study, no change in perception was observed. Furthermore, Torta et al. (2017) showed an enhancement of the vertex N wave elicited by visual stimuli projected onto the arm at which HFS was delivered. The visual stimuli were generated by a visible laser beam at a wavelength clearly inadequate to activate heat and mechanoreceptors in the skin. Therefore, it seems that HFS can induce effects that are not restricted to the transmission of nociceptive input at the level of the spinal cord.

However, the enhancement of the N wave elicited by the visual stimuli was short-lasting, being present at 20 minutes but not 45 minutes after applying HFS, which is in contrast to the long-lasting enhancement observed for vibrotactile stimuli and IES which is present both 20 minutes and 45 minutes after applying HFS. A possibility could be that these differences in time-courses are related to the modality of the stimulus. Somatosensory stimuli applied onto the arm at which cutaneous hyperalgesia is induced may be more salient or relevant than visual stimuli.

To conclude, HFS induced a long-lasting enhancement of the N2 wave elicited by IES in the area of secondary hyperalgesia, indicating that HFS enhances the responsiveness of the central nervous system to nociceptive inputs conveyed by AMH-II nociceptors. However, we found no evidence that HFS affects the perception elicited by IES, which may suggest that AMH-II nociceptors do not contribute to HFS-induced secondary hyperalgesia.

## ACKNOWLEDGEMENT

The authors would like to thank Federico Arguissain and Fabricio Ariel Jure for their help.

## AUTHOR CONTRIBUTIONS

Conception and design: EvdB, AM, JBM. Acquisition of data: EvdB, JBM. Analysis and interpretation of data: EvdB, AM, JBM, OKA. Drafting the manuscript: EvdB, AM, JBM, OKA. Final approval of manuscript: EvdB, AM, JBM, OKA.

