## Supplementary Materials for "High frequency electrical stimulation induces a long-lasting enhancement of event-related potentials but does not change the perception elicited by intra-epidermal electrical stimuli delivered to the area of secondary mechanical hyperalgesia"

### Supporting information

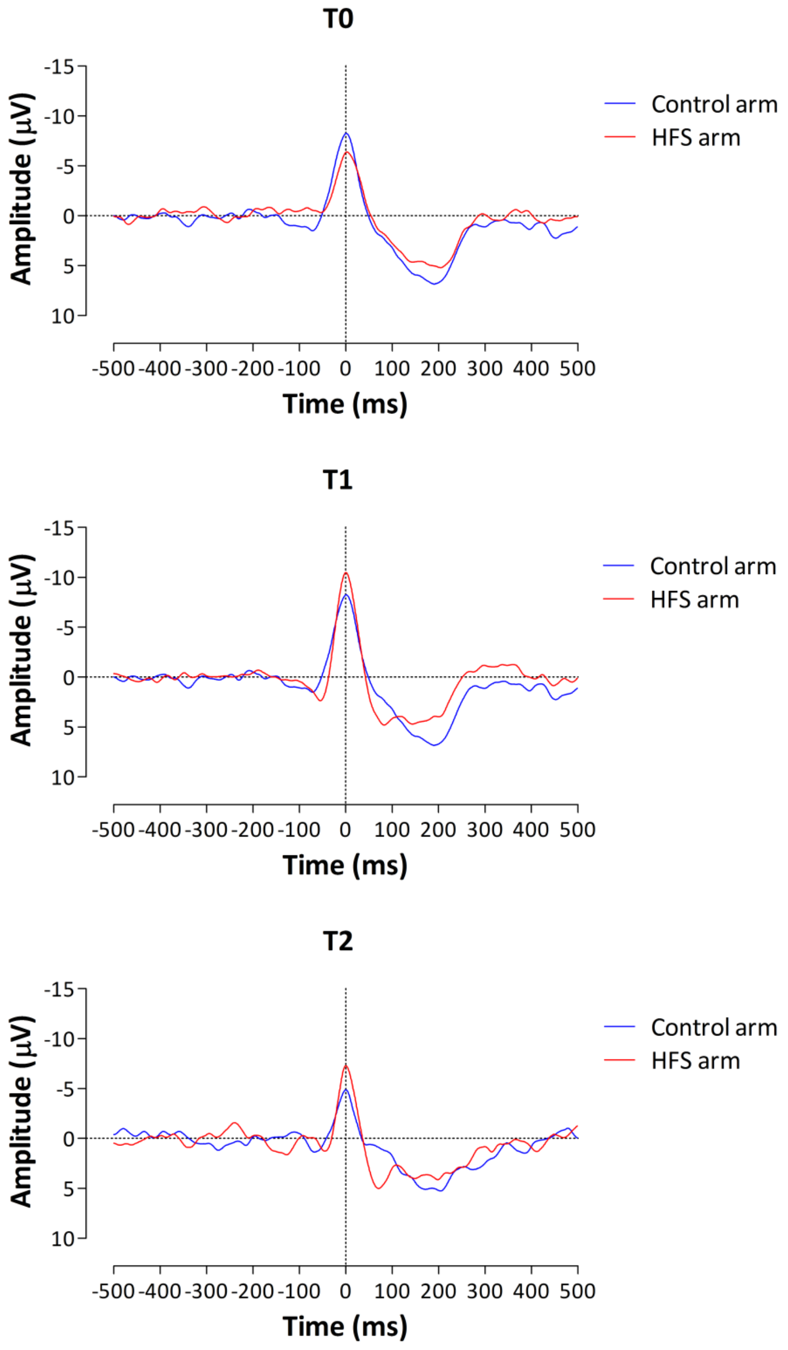


**Figure S1.** Aligned (to the individual N2 peaks) group-level average event-related potentials (Cz) elicited by intra-epidermal electrical stimulation before and after applying HFS from both arms.
